# Content-Sensitive Linguistic Representations in the Human Multiple-Demand Network

**DOI:** 10.64898/2026.09.14.749455

**Authors:** Miriam Havin, Meir Meshulam, Taelin Karidi, Refael Tikochinski, Uri Hasson, Ariel Goldstein

## Abstract

Neuroimaging dissociates specialized language regions from the domain-general multiple-demand (MD) network, yet the functional contribution of MD regions to language processing remains unresolved. Because MD recruitment during linguistic tasks is conventionally attributed to domain-general cognitive load, prior research has largely prioritized activation magnitude over representational content. Consequently, it remains unclear whether MD cortices function merely as nonspecific amplifiers of effort or actively encode fine-grained linguistic structures.

Here we show that the MD network reliably encodes content-sensitive linguistic representations during naturalistic comprehension independently of peak cognitive demand. Using a voxelwise encoding framework with architecturally identical language models trained on distinct semantic domains (BERT and SciBERT) across three naturalistic fMRI datasets, we find that multiple-demand regions track linguistic structure during both high-demand scientific lectures and low-demand narratives. Furthermore, comparative analyses reveal that these representations are sensitive to domain-specific semantic regularities rather than surface-level form alone. Finally, by evaluating relative model alignment alongside the language network, we demonstrate that the representational balance between these two systems shifts dynamically as a function of task context and domain relevance, revealing a complementary division of labor. Together, these findings reframe the multiple-demand network from a purely extrinsic control mechanism into an active, content-sensitive component of a distributed semantic architecture.

## Introduction

Functional neuroimaging research has traditionally characterized the human brain through a lens of strict network specialization. In particular, the multiple-demand (MD) network has been linked to domain-general executive control (Duncan, 2010; Duncan & Owen, 2000; Duncan, 2013), while the language network has been associated with linguistic processing (Fedorenko et al., 2011; Fedorenko et al., 2012). This dissociation has motivated a widely held view of functional specialization, in which language comprehension is primarily supported by a dedicated language-selective network, whereas the MD network contributes domain-general executive processes that are recruited only when comprehension imposes additional cognitive demands, such as increased working memory load, ambiguity resolution, or extraneous task demands, such as a picture naming task or a semantic judgment task (Billot et al., 2026, Fedorenko et al., 2014; Fedorenko & Blank, 2020).

A recurring observation in the literature is that the MD network is frequently recruited alongside the language network during complex language tasks (Fedorenko et al., 2013; Blank et al., 2014; Wehbe et al., 2020; Diachek et al., 2020). This recruitment has been observed across a range of paradigms, particularly those involving explicit task demands. Because MD activations vary systematically with task design, they have generally been interpreted within a domain-general framework, in which MD activity reflects executive processes engaged by task demands rather than language-specific computations (Blank et al., 2014; Diachek et al., 2020; Müller et al., 2024). However, this interpretation has been strongly informed by analyses of activation magnitude in controlled experimental paradigms, where task and stimulus contributions can be more cleanly dissociated (Blank et al., 2014; Fedorenko et al., 2011; Diachek et al., 2020). While this approach has been instrumental in establishing the functional dissociation between the language and MD networks, activation magnitude alone cannot determine if MD responses reflect task demands or content-dependent representations. Consequently, these studies provide limited insight into the representational structure encoded within the MD network during language comprehension. As a result, the MD network has been characterized primarily in terms of when it is engaged, rather than what it represents.

More generally, the nature of representations within the MD network remains an open question. One hypothesis is that it supports task performance by providing cognitive control and coordination across specialized systems (Duncan, 2010; Cole et al., 2013). An alternative hypothesis is that, in addition to that role, MD regions directly encode and process task-relevant representations in concert with specialized regions (Woolgar et al., 2011; Woolgar et al., 2016; Shashidhara et al., 2021). Language comprehension provides a useful lens through which we can address this question. If MD activity exclusively reflects executive demands, one would expect little systematic encoding of linguistic content in MD regions. Conversely, finding evidence of linguistic processing in MD activity on par with language regions would suggest that the MD network participates more directly in representing information during ongoing cognition.

Moreover, as recent evidence demonstrates that rich linguistic and semantic representations extend far beyond the core language-selective cortex, a deeper theoretical challenge emerges (Huth et al., 2016; Goldstein et al., 2022; Ivanova et al., 2025; Federenko et al., 2025). When powerful encoding models and naturalistic paradigms reveal that rich linguistic and semantic information can be decoded widely across the cortex, it forces us to re-evaluate the foundational concept of network specialization itself. Does widespread, distributed representation imply that functional boundaries have collapsed into undifferentiated processing, or can we still articulate a nuanced, mechanistically distinct view of different networks within a whole-brain representational account? Within this landscape, we investigate the multiple-demand network not merely to categorize its response to task load, but to test how a domain-general executive system maintains functional differentiation while tracking the complex semantic structure of naturalistic experience.

To resolve this tension, we investigate the representational sensitivity of the multiple-demand network during naturalistic language comprehension. Rather than viewing distributed representations as evidence of functional collapse, we test how a domain-general executive system maintains mechanistic differentiation from the language network while both are co-activated. Our analysis proceeds in three stages: first, we determine whether the MD network encodes linguistic information; second, we test whether these representations are sensitive to semantic structure or merely to surface-level linguistic form; and third, we compare this semantic sensitivity against that of the language network to determine whether distinct functional profiles persist within a shared, distributed informational space.

## Methods

### Participants and Datasets

Our study utilized fMRI data from three independent cohorts to investigate the neural alignment of domain-adapted language models.

#### Dataset 1 (Computer Science Lectures)

This dataset is a reanalysis of a large-scale naturalistic fMRI study of learning and comprehension (Meshulam et al., 2021). It consists of N=24 undergraduate participants scanned over a 13-week semester while viewing 21 segments of computer science lecture videos (total duration: 197 minutes). All participants had no prior formal knowledge of the material.

#### Dataset 2 (Narratives)

Dataset 2 consisted of an analysis of a naturalistic auditory narrative fMRI dataset comprising continuous story listening data from *The Moth Radio Hour* and a complementary publicly available naturalistic language comprehension dataset (Huth et al., 2016). The dataset includes N=9 participants who listened to approximately 370 minutes of continuous spoken narrative per participant.

#### Dataset 3 (Single-Video computer science lecture and narrative)

To provide an additional within-subject comparison, we analyzed data from an independent cohort of N=21 participants. Each participant viewed two single-video stimuli: one computer science lecture (duration: 7 minutes) and one story from *The Moth* podcast (duration: 13 minutes). This dataset was used to isolate stimulus-driven differences in network engagement and to verify that effects observed in the full datasets generalize to brief, single-episode presentations.

### Neural Data Acquisition and Preprocessing

For Dataset 1, functional images were acquired with a repetition time (TR) of 2000∼ms and 3∼mm isotropic voxels. For Dataset 2, functional images were acquired with a TR of 2000ms and 3mm isotropic voxels. For Dataset 3, images were acquired with a TR of 1500ms and 2mm isotropic voxels. All datasets underwent standard preprocessing procedures including motion correction and slice-timing correction. Dataset 1 and Dataset 3 were additionally aligned to the MNI-152 anatomical template, whereas Dataset 2 was analyzed in native functional space following the preprocessing and alignment procedures of the original pipelines. Full acquisition parameters and preprocessing details for Datasets 1 and 2 are reported in the original publications (Meshulam et al., 2021; Huth et al., 2016), and the same preprocessing pipeline as dataset 1 was applied to Dataset 3.

### Network Definitions

#### Multiple-Demand (MD) Network

The multiple-demand (MD) network was defined using a probabilistic atlas derived from functional localizer data collected in approximately 200 participants (Fedorenko et al., 2013; Assem et al., 2020). These localizers identify regions that respond more strongly to cognitively demanding than to less demanding conditions across a variety of tasks. The resulting network comprises bilateral frontal and parietal regions consistently engaged across task domains, including lateral prefrontal cortex, anterior insula, pre-supplementary motor area, and intraparietal sulcus. The MD network has been implicated in domain-general cognitive control and flexible information integration.

#### Language Network

The language network was defined using a probabilistic atlas constructed from sentence-processing localizer data collected in approximately 200 participants (Fedorenko et al., 2010; Fedorenko et al., 2011). These localizers identify regions that respond more strongly to meaningful sentences than to nonword or word-list control stimuli. The resulting network includes predominantly left-lateralized frontal and temporal regions selectively engaged during high-level linguistic processing, including portions of the inferior frontal gyrus and lateral temporal cortex. These regions are selectively responsive to linguistic structure and meaning and are functionally dissociated from domain-general control systems.

#### Atlas Application

All network masks were defined independently of the present data and analyses and were applied as fixed regions of interest.

### Computational Language Models

#### Architectures and Weights

We compared two transformer-based language models: a general-purpose model (BERT) and a domain-adapted model (SciBERT). Both models are based on the BERT-base architecture (Devlin et al., 2019), consisting of 12 transformer layers with 768-dimensional hidden representations. Critically, we used versions of both models that share an identical WordPiece vocabulary (BaseVocab). By holding the architecture and tokenization scheme constant, any differences in neural predictive power can be attributed specifically to differences in the learned model weights rather than to differences in token segmentation.

#### Pre-training

BERT (Base, Uncased) was pre-trained on the BooksCorpus and English Wikipedia, reflecting general-domain English usage. SciBERT was pre-trained on a corpus of 1.14 million scientific papers from Semantic Scholar (Beltagy et al., 2019). Although the two models share the same architecture and vocabulary, SciBERT’s parameters were optimized on scientific text, enabling it to capture structural and conceptual regularities characteristic of scientific discourse.

### Feature Extraction and Encoding Framework

#### Feature Extraction and Alignment

To relate language model representations to neural activity, we extracted contextualized embeddings from BERT and SciBERT using a sliding context window approach. For each word in the transcript, a context window of 32 tokens was constructed and passed through the model. Following prior work, we used a 32-token context window, which has been shown to yield maximal neural alignment in voxelwise encoding analyses, with longer contexts providing no additional benefit (Tikochinski et al., 2025). Token level hidden state representations were averaged to obtain a single word-level embedding. Embeddings were extracted from all 12 transformer layers.

Word embeddings were temporally aligned to the fMRI time series using Lanczos interpolation, a band-limited resampling method previously used to align continuous stimulus features to discrete fMRI sampling times (Huth et al., 2016). To account for the temporal dynamics of the hemodynamic response, stimulus feature matrices were expanded to include four delayed copies of each feature corresponding to successive post-stimulus time lags spanning from 2 to 8 seconds. These delayed features were concatenated and used as predictors in the encoding models, a procedure standard for voxelwise encoding models (Huth et al., 2016). Unless otherwise specified, primary model comparisons were conducted using representations from the final transformer layer.

**Fig 1.**
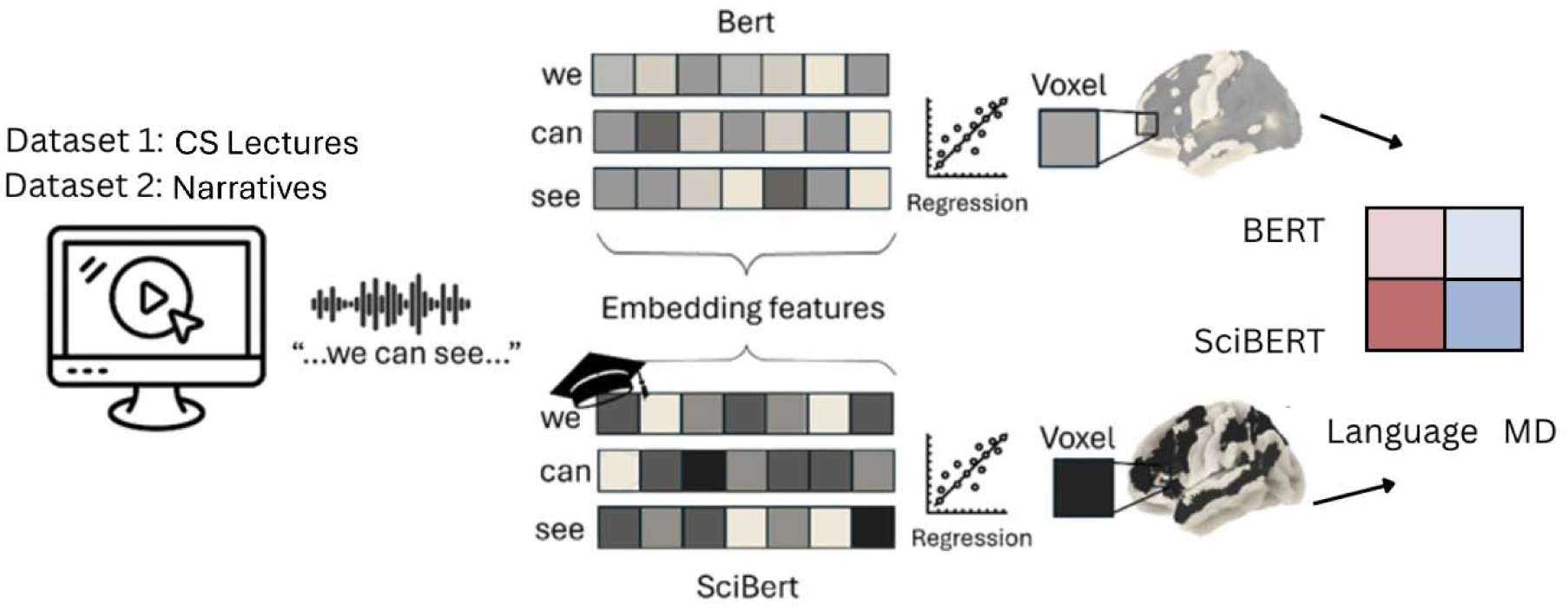
Computational encoding pipeline and comparative framework. Contextual representations from SciBERT and BERT are extracted from lecture transcripts and used to predict voxelwise brain responses during lecture viewing. Model differences are assessed by computing voxelwise predictions from each language model (BERT and SciBERT), which are then summarized across the cortex and within predefined multiple-demand and language networks.

#### Encoding Model

Neural responses were modeled using a voxelwise linear encoding framework. Models were fitted independently for each voxel using ridge regression. The regularization parameter alpha was selected from a logarithmically spaced range using cross-validation. Encoding performance was evaluated using five-fold cross-validation. To reduce variance in hyperparameter selection under naturalistic noise, ridge regression within each training fold was repeated across 50 bootstrap resamples, and alpha was chosen based on average cross-validated performance across resamples.

#### Evaluation and Voxel Selection

Encoding performance was quantified as the Pearson correlation between predicted and observed BOLD responses on held-out data, with all correlation values Fisher z-transformed prior to group-level analyses. Gains were averaged within predefined Language and Multiple-Demand network parcels then compared across participants.

## Results

In this paper we present evidence that the MD network stores linguistic information during passive comprehension regardless of the level of cognitive demand. We further show that the information represented by the MD network is specifically sensitive to semantic content. Finally, we find that this sensitivity to content characterizes the MD network more than the language network, providing a potential differentiation between the two networks, despite both encoding linguistic information during passive listening.

In order to support these claims, we first verify the two datasets indeed differ in their levels of cognitive demand. We then test whether the MD network carries linguistic information by linearly mapping LLM based representations to the brain signal. Next, we test whether the network shows sensitivity to semantic content by comparing mappings from models with different training data. Finally, we examine the interaction between the different models across networks, in order to characterize the sensitivity of each network to model differences.

Throughout the analyses, we treat the language network as a validation network. Given its established sensitivity to linguistic structure, it serves as a reference system for validating the framework and for interpreting MD network responses. All analyses applied to the Language network are identical to those applied to the MD network, allowing direct comparison across systems.

### Manipulation Check on cognitive demand

A central aim of this study is to analyze how the MD network represents language. Given that this network is characterized by its responsiveness to cognitive demand, we examine its linguistic representations across a spectrum of difficulty. We utilize two datasets that span this spectrum: a high-demand introductory computer science course (24 students) characterized by conceptual abstraction, and a lower-demand autobiographical narrative from ‘The Moth’ podcast (8 subjects) characterized by narrative structure and lower information density. We hypothesize that these datasets reliably differ in their imposed processing demands, providing a framework to test whether the MD network’s linguistic sensitivity persists independently of task effort.

Since the two datasets were independently collected and therefore differ along multiple dimensions including subjects, acquisition parameters and stimuli, differences between them cannot reliably be attributed to stimuli alone. Therefore, we utilized a third dataset with a within-subject experiment design, where subjects watched one 10 minute long video of the computer science course and one 10 minute long video of the stories. Although too short to support the full set of analyses in this study, this dataset provides a controlled setting, allowing us to isolate the effect of stimulus and validate differences in MD activation across conditions.

We confirmed the difference in cognitive demand through a General Linear Model (GLM) analysis, which yielded a robust interaction effect between brain networks (Language vs. MD) and stimulus types (t(20) = −2.65, p = 0.016, d = −0.58; Fig. 2). This interaction indicates that the effects of stimulus type differ across the Language and MD networks. Overall, the MD network shows higher sensitivity to the computer science lecture relative to the Language network, consistent with the expected difference in cognitive demand between the two stimulus types.

**Fig. 2.**
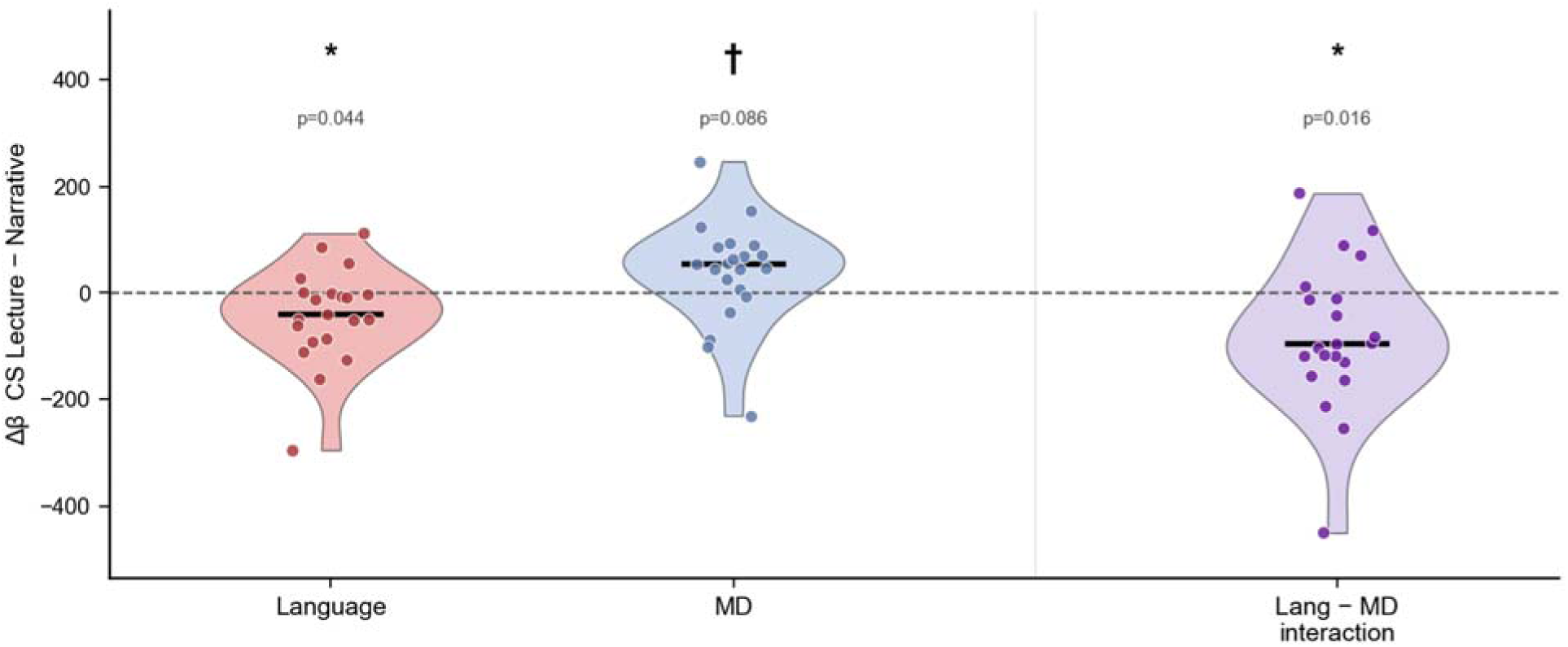
MD network engagement is selectively modulated by stimulus content. Mean engagement of Language and MD networks during high and low demand tasks (N = 21). A robust interaction (t(20) = -2.645, p = 0.016) reveals that the MD network shows higher recruitment than the language network during the Computer Science lecture, indicating increased executive demand. Error bars reflect SEM.

### Neural Encoding of Linguistic Information in Language and MD networks

Having verified that the two datasets differ in MD response magnitude, we move to explore the amount of linguistic information in the MD network stored in each dataset. While the Language network is the canonical system dedicated to linguistic processing, we hypothesized that if the MD network contributed to high-level understanding during naturalistic linguistic tasks, it should also maintain a linguistic representation, even when not explicitly required to perform a task.

To do this, we employed encoding models. These models use prediction as a proxy for neural representation: if a model can accurately predict the fluctuations in the BOLD signal using specific linguistic features, it suggests that the brain is using those same features to process the information. By mapping these features onto neural activity through voxel-wise linear regression, we quantify the extent to which model-derived linguistic features account for variance in each voxel’s response. This approach allows us to isolate and compare the information structure processed by the Language and MD networks.

Using a pre-trained BERT model, we generated embedding representations for the stimuli from both datasets. These embeddings were aligned to the neural time series and used as predictors in the encoding models. We then averaged the encoding score per voxel over each network’s atlas, yielding a single network-level encoding score per dataset and participant. We first applied this framework to the Language network as a validation of the analysis approach, given its established sensitivity to linguistic structure. Encoding performance in this network was robustly above chance (flipped sign permutation test, high demand, p < 0.001, low demand p=0.001; Fig. 3A), confirming that the model captures expected language-related neural structure.

**Fig. 3.**
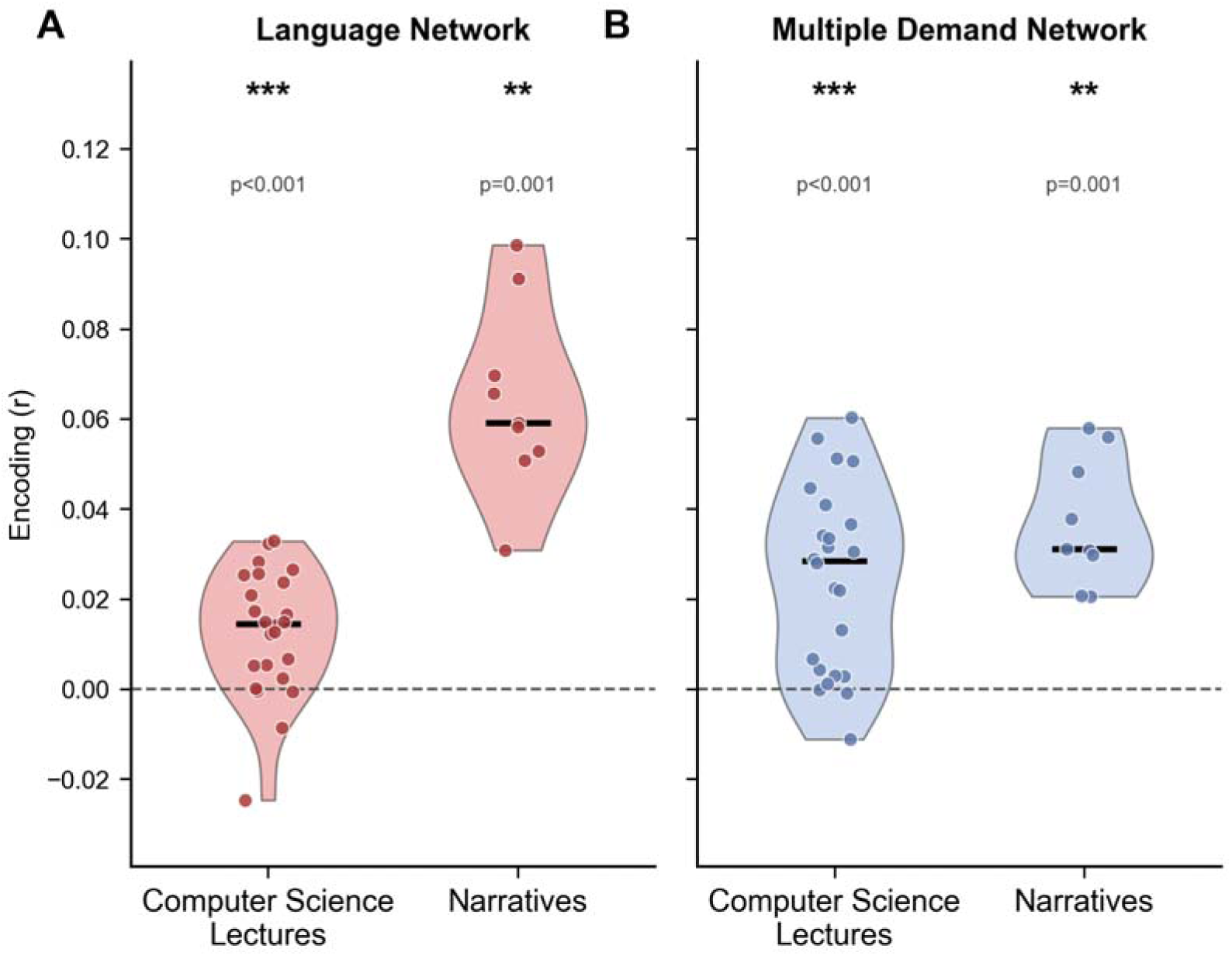
Language and Multiple-Demand networks exhibit robust, significant linguistic encoding accuracy across high- and low-demand naturalistic datasets. (A) Language network and (B) Multiple-Demand (MD) network encoding accuracy (Fisher z) for the BERT model across high and low demand datasets. Both networks exhibit significant encoding performance in both domains. Flipped sign permutation tests confirm significance for all conditions: high demand (p < 0.001) and low demand (p = 0.001).

We then applied the same encoding framework to the MD network. Encoding performance in the MD network was significantly above chance in both datasets (flipped sign permutation test, high demand, p < 0.001, low demand p=0.001; Fig. 3B), indicating that linguistic information is reliably encoded in MD regions during passive comprehension regardless of cognitive demand. Full brain encoding plots can be seen in supplementary figure 1.

Additionally, we found that the two networks exhibit distinct profiles of information tracking (Fig. 3). In the high demand task, the MD network demonstrated significantly higher encoding accuracy than the Language network (BERT: p=0.0002; SciBERT: p<0.0001). Conversely, in the low demand task, this pattern was significantly reversed, with the Language network showing stronger tracking fidelity (BERT: p<0.0001; SciBERT: p<0.0001). This interaction was robustly replicated across both models (Interaction: p<0.0001).

These results largely recapitulate the previously reported pattern of network recruitment. Consistent with prior activation-based findings, the MD network showed stronger encoding during the more demanding computer science lecture, whereas the Language network showed stronger encoding during the narrative stimulus. This correspondence suggests that the cognitive factors associated with differential network recruitment also influence representational encoding. However, differences in encoding strength may partly reflect changes in signal magnitude or reliability, and therefore do not necessarily indicate that one context elicits more informative representations than another. Instead, they suggest that network engagement and representational strength are jointly shaped by cognitive demands.

### Content-Sensitive Modulation in Language and MD Networks

Having established that the MD network represents linguistic information, we next investigated whether this representation is specifically sensitive to the semantic content of the discourse, rather than the linguistic form. To do this, we compared neural alignment between two language models: BERT, trained on general-purpose corpora, and SciBERT, which shares the same architecture but is trained on scientific publications. Because these models differ only in their training data, any divergence in neural encoding provides a controlled measure of how domain-specific training distributions map onto the brain.

In the Language network, we observed consistent sensitivity to differences between models, reflected in a significant interaction between model type and stimulus Interaction: t(31) = -2.406, p = 0.022) (Fig. 4A). SciBERT shows comparable brain alignment to BERT for high demand dataset (p = 0.052), but lower alignment for low demand dataset (p = 0.0044).

**Fig. 4.**
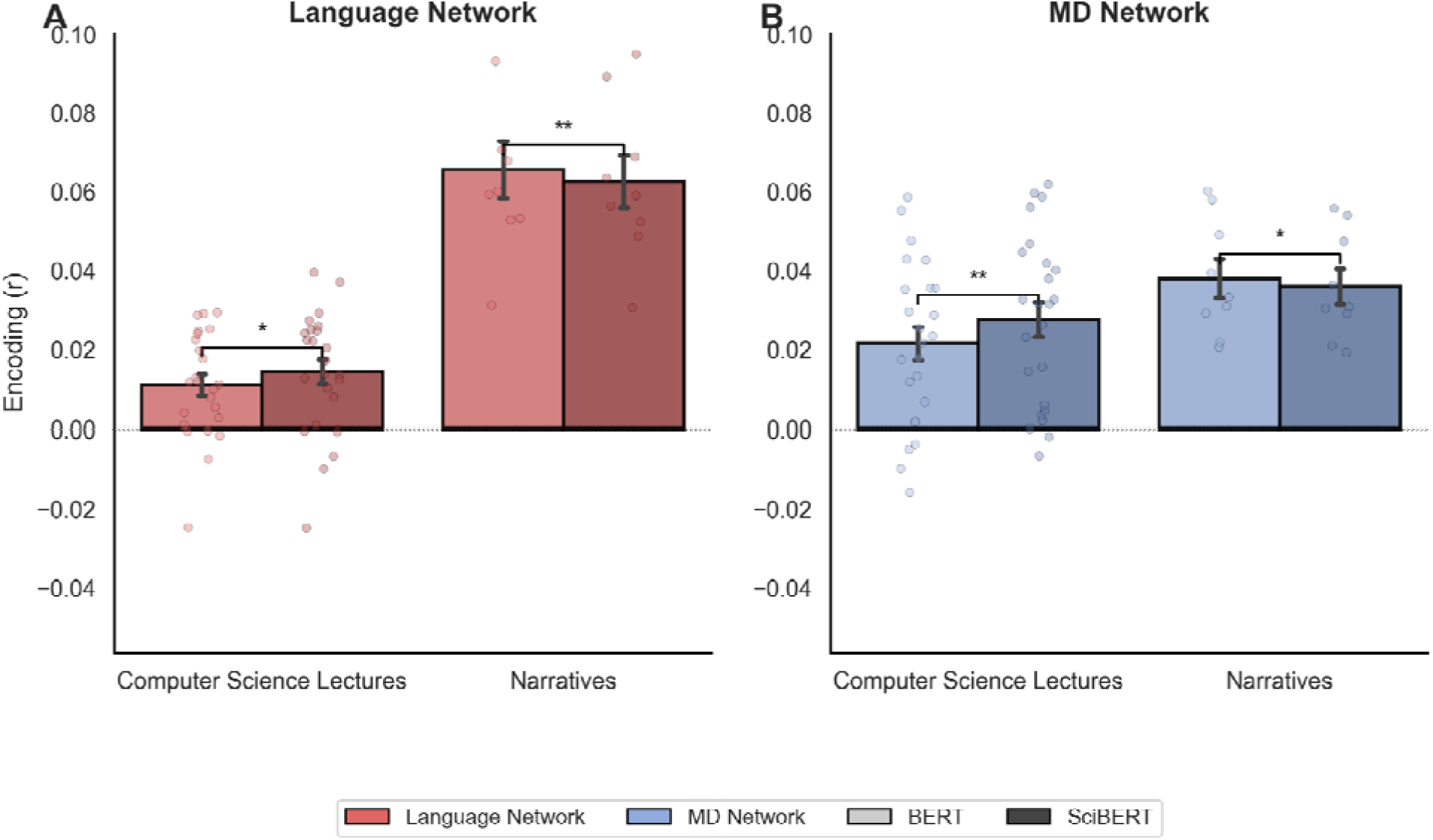
Differential model alignment reflects content-specific semantic sensitivity across networks. Encoding (Fisher z) across Language and MD networks. (A) Language network: SciBERT matches BERT on scientific content (p = 0.052) but drops on narratives (p = 0.0044). (B) MD network: SciBERT exceeds BERT on lectures (p = 0.0058), while BERT exceeds SciBERT on narratives (p = 0.0192). N=24 (Computer Science Lectures) and N=9 (Narratives).

This indicates that model-derived representations differ in how well they predict neural responses depending on the training corpus. Rather than one model being universally better, each model provides a feature space that aligns more closely with brain activity when the input matches its training distribution. SciBERT’s representations therefore support similar levels of neural predictability as BERT in scientific contexts, but lose alignment when the discourse shifts outside that domain, even if it remains coherent and information-rich.

Once establishing that the difference between models captures content differences in the language network, we investigated whether the MD network displays similar linguistic tracking. As demonstrated in Fig 4. B, the MD network replicates the linguistic tracking observed in the Language network, confirming that it tracks linguistic features across both the high and low demand datasets.

Crucially, the MD network exhibits content sensitivity in both datasets: it demonstrates robust tracking of linguistic information during the high-demand discourse (p = 0.0044), yet maintains a distinct, reliable sensitivity to linguistic features even within the low-demand dataset (p = 0.0192). This indicates that the MD network’s linguistic representation is not merely an artifact of peak cognitive effort; rather, it is modulated by the context of the stimulus across the entire range of difficulty. These findings demonstrate that the MD network provides a content-sensitive contextualization of the linguistic structure.

### Training-Driven Modulation of Network Selectivity

Having established that both networks track linguistic information and are sensitive to content, we sought to determine if they are functionally specialized. Simply showing that both networks respond to semantic content does not reveal whether they process that content identically or if they serve distinct roles. To explore this we leveraged the fact that SciBERT and BERT are architecturally identical but capture different semantic domains. By evaluating how relative neural alignment shifts as a function of training-content match, we can isolate whether the MD network is differentially tuned to semantic features driven by domain-relevant training.

To quantify this, we computed a Differential Sensitivity Index for each subject, capturing the relative representational bias between the MD and Language networks. We found that this index varied systematically by task context: the interaction was significantly positive in the high-demand scientific dataset (t(23) = 2.12, p = 0.045), reflecting a greater sensitivity to domain-specific training differences within the MD network. Conversely, the effect reversed in the low-demand narrative dataset (t(8) = 2.46, p = 0.040), where the Language network showed greater sensitivity to these model differences (see Fig. 5).

**Fig. 5.**
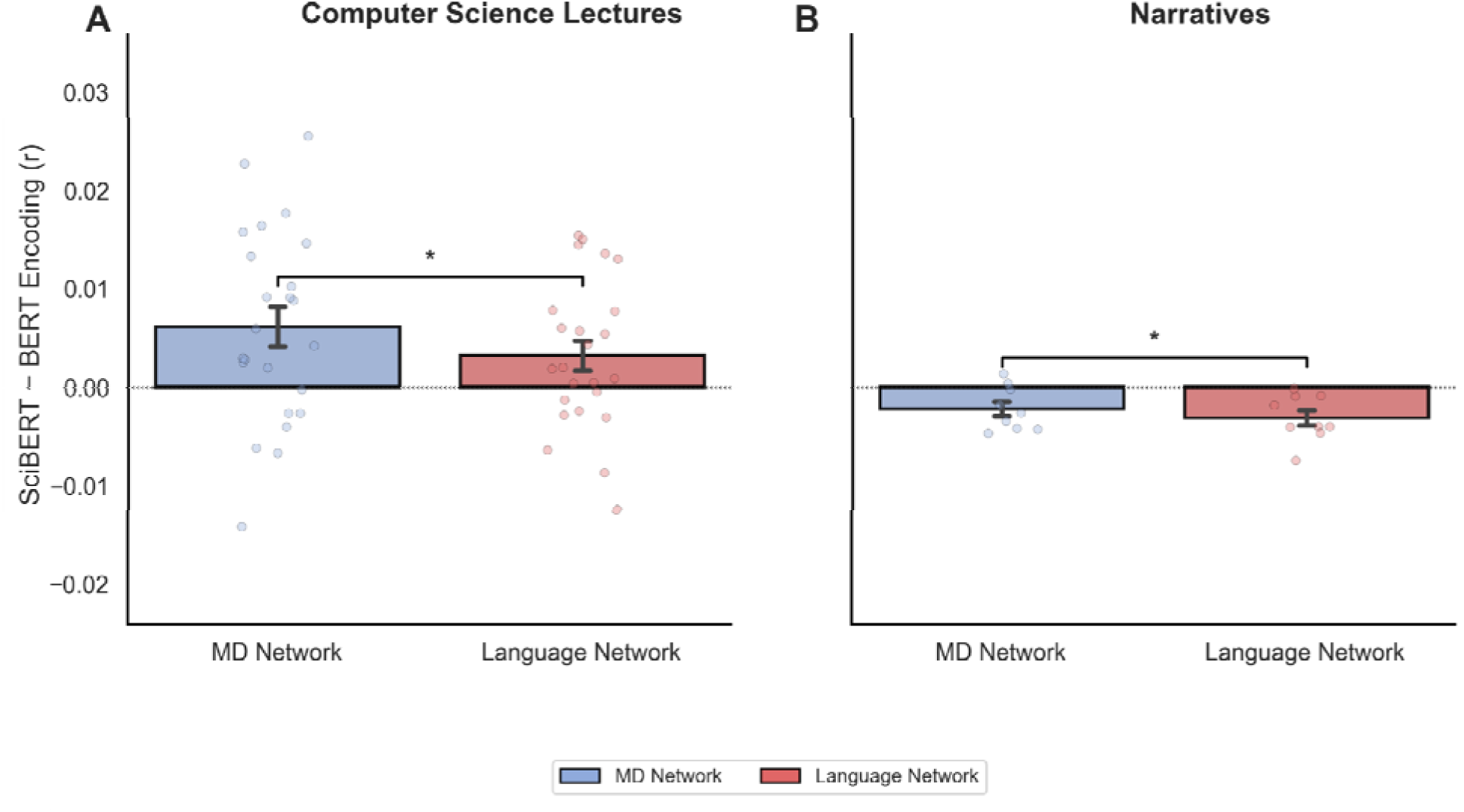
Differential model alignment showing that task context shifts the relative representational balance between the MD and Language networks. Difference in encoding accuracy (SciBERT - BERT, Fisher z) between MD (blue) and Language (red) networks for high demand (Computer Science Lecture) (A) and low demand (Narrative) (B) tasks. Positive values indicate higher alignment for content relevant models, showing that the relative representational balance shifts toward the MD network during scientific discourse and toward the Language network during narrative processing. Error bars: SEM; *: p < 0.05.

These results indicate that while both networks encode linguistic information, they are differentially tuned depending on task context. The MD network exhibits heightened sensitivity to domain-specific training differences during scientific discourse, whereas the Language network shows greater sensitivity during narrative comprehension, suggesting that these systems dynamically adjust their representational balance to complement each other across varying cognitive demands.

## Discussion

The present study aimed to re-evaluate the processing of the Multiple-Demand (MD) network in language comprehension by moving beyond traditional activation-based metrics. By employing a voxelwise encoding framework and comparing neural responses to architecturally identical language models with divergent training histories, we provide evidence that the MD network does more than simply respond to the "extrinsic" cognitive demands of a task.

Our findings support three primary conclusions: first, the MD network reliably encodes information captured by linguistic model representations during naturalistic comprehension, including during relatively low-demand contexts; second, these representations are sensitive to domain-relevant statistical regularities rather than surface-level linguistic features alone, as suggested by the differences between models trained on distinct semantic domains; and third, the MD network displays a distinct representational signature compared to the Language network, characterized by a unique profile of model sensitivity that varies across experimental contexts. Collectively, these results demonstrate that the relative contribution of the MD and Language networks to semantic representation shifts dynamically as a function of domain-relevance and cognitive context, revealing a complementary division of labor rather than a static sharing of linguistic content.

### Reframing MD Network Functionality

Our findings contribute to an emerging literature that shifts the characterization of the MD network away from a singular domain-general control mechanism toward a system capable of representing task-relevant information (Cole et al., 2013; Woolgar et al., 2016; Assem et al., 2020). Historically, MD recruitment during language tasks has been interpreted as reflecting domain-general executive demands that emerge when linguistic processing becomes effortful or when additional cognitive control is required (Fedorenko et al., 2013; Blank et al., 2014). These activation-based studies have been instrumental in defining the functional dissociation between the Language and MD networks. However, univariate activation magnitude provides limited insight into the representational content carried by these responses. By moving beyond activation to a representational account, our results suggest that co-activation of Language and MD networks during comprehension may reflect parallel streams of information processing rather than a simple division between linguistic computation and domain-general effort.

Several aspects of the present results support this reinterpretation of MD function. First, the ability of language model representations to predict MD responses indicates that these regions carry information aligned with linguistic representations, rather than reflecting only nonspecific cognitive demands. Moreover, differences in prediction accuracy between architecturally identical models trained on distinct semantic corpora suggest that this information is sensitive to semantic content rather than generic linguistic features. Second, the relative sensitivity of the MD and Language networks varied across datasets, indicating that these systems make flexible and context-dependent contributions to semantic representation rather than reflecting a fixed division between linguistic computation and domain-general control.

### Distributed Representations Across Language and Association Cortex

This emerging view of MD functionality aligns with broader shifts in cognitive neuroscience away from strictly modular accounts of cognition toward models in which information is represented across distributed, interacting cortical systems. Recent work using naturalistic stimuli and computational models has demonstrated that semantic information is not confined to classical language regions but is distributed throughout association cortex, including regions traditionally associated with executive and control functions (Huth et al., 2016; Pereira et al., 2018; Caucheteux & King, 2022).

Within this framework, the present findings suggest that the MD network may represent one component of a broader architecture in which information derived from linguistic input is distributed across multiple cortical systems. Rather than indicating that MD constitutes an additional language system, these findings suggest that comprehension engages multiple interacting representational systems, with MD contributing information that complements the specialized linguistic computations supported by the Language network.

One possibility is that Language regions primarily maintain representations optimized for extracting and manipulating linguistic structure, whereas MD regions integrate these representations with broader contextual, goal-related, and domain-general information required for flexible behavior. This integration may be selectively engaged when comprehension places greater demands on maintaining, manipulating, or coordinating information, allowing MD contributions to increase when additional cognitive control is required. Thus, MD representations may reflect a flexible resource that is recruited according to the demands of the current context rather than a fixed component of all language processing.

### Breaking the Co-activation Barrier

A significant challenge in neuroimaging research is disentangling representational content from the cognitive operations required to process that content. Traditional paradigms relying on activation magnitude can identify networks recruited during a task but cannot determine whether jointly activated regions represent similar information or participate in distinct computational processes.

Our computational framework addresses this limitation by leveraging language models as representational probes. By holding model architecture and tokenization constant while manipulating training history, we created a controlled approach for testing whether neural responses preferentially align with specific learned statistical structures. This approach builds upon a rapidly expanding literature using artificial neural networks to investigate the computational principles underlying human cognition (Kriegeskorte et al., 2018; Yamins & DiCarlo, 2016; Goldstein, 2022). Rather than treating language models merely as predictive tools, recent approaches increasingly use them as hypotheses about the representational spaces that may support biological cognition (Friston et al., 2009).

### Limitations and Robustness

While our findings provide a novel perspective on the MD network, they should be interpreted within the constraints of our experimental design. The primary limitation arises from dataset heterogeneity: the high-demand (Computer Science lecture) and low-demand (narrative) datasets were collected from independent cohorts with different acquisition parameters and preprocessing pipelines, which could introduce systematic variance. However, the fact that MD sensitivity was observed despite these differences, and was further validated in a within-subject control dataset with matched acquisition and preprocessing, suggests that the observed effects are unlikely to be explained solely by dataset-specific factors. Future studies using larger, unified cohorts will be important for establishing the stability and generalizability of these representational patterns.

A further consideration is that the MD network is not a homogeneous system. Recent work suggests functional and anatomical heterogeneity among MD components, with different regions contributing to distinct aspects of cognitive control, abstraction, and representation (Duncan, 2013; Assem et al., 2020). Because the present analysis examined network-level effects, future studies using finer-grained parcellations or individual-subject functional localization will be necessary to determine whether representational sensitivity is distributed across MD or concentrated within specific subregions.

Finally, although the comparison between BERT and SciBERT provides a controlled manipulation of model training history, it does not isolate semantic knowledge from other properties of scientific language, including lexical statistics, discourse structure, and domain-specific regularities. Future work using more targeted model comparisons will be necessary to identify the specific representational features underlying MD sensitivity. BERT and SciBERT were selected because their shared architecture and divergent training corpora allow differences in neural sensitivity to be attributed more directly to learned representations rather than architectural differences. However, an important direction for future work will be to determine whether these findings generalize across newer language models with different architectures, scales, and training objectives.

### Future Directions

Our findings suggest several directions for future investigation. First, rather than viewing Language and MD networks as representing a binary distinction between specialized linguistic processing and general cognitive control, future work should examine how these systems dynamically interact during comprehension. A central question is whether MD involvement increases during moments requiring maintenance of broader conceptual context, integration across longer timescales, or resolution of semantic uncertainty.

Second, the present findings raise questions about how representational sensitivity within MD emerges through experience. If MD regions contribute to context-dependent representations of learned knowledge, then expertise and training may reshape the information represented within these regions. This possibility connects with broader theories of neural plasticity in which cortical representations are continuously adapted to reflect the demands of an individual’s environment and accumulated knowledge.

Finally, continued integration between neuroscience and artificial intelligence provides new opportunities for understanding the computational organization of cognition. As increasingly powerful models allow more precise manipulation of representational structure, future studies can move beyond correlational alignment toward testing whether specific computational properties of artificial systems predict corresponding changes in biological representations.

## Acknowledgements

We would like to thank Timna Wharton Kleinman and Daria Lioubashevsky for their helpful feedback and valuable insights.

This work was supported by the European Research Council (ERC) under the European Union’s Horizon Europe research and innovation programme (ERC Starting Grant “WordOrigions”, Grant Agreement No. 101222215).

## Supplementary Figures

**Fig. S1.**
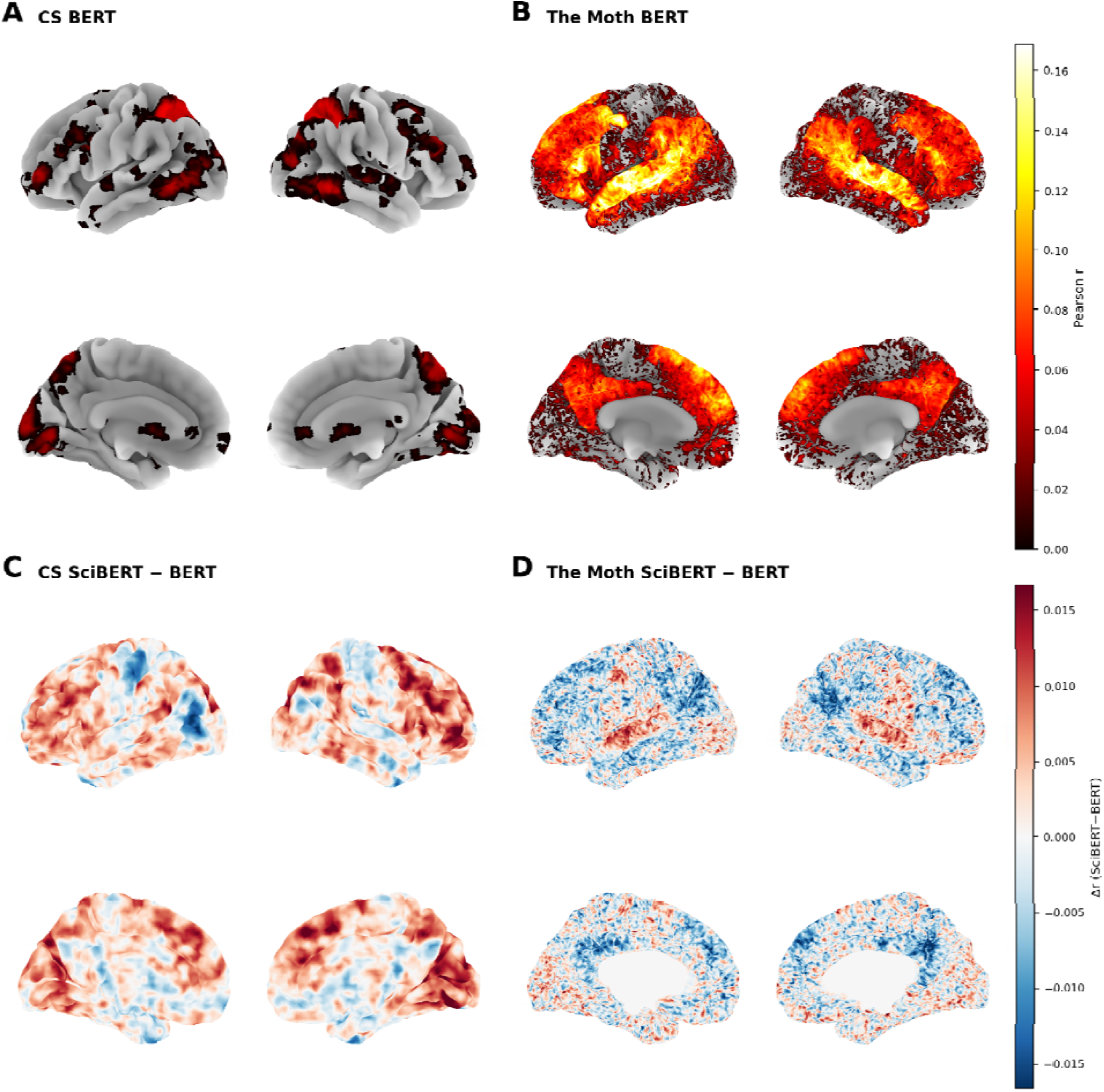
Voxelwise encoding accuracy maps for BERT and difference maps between SciBERT and BERT. Encoding performance (Pearson r) for the BERT model during (A) Computer Science lectures and (B) The Moth narratives. Difference maps (Δ r, SciBERT minus BERT) for (C) Computer Science lectures and (D) The Moth narratives.

## Notes

### Competing Interest Statement

The authors have declared no competing interest.

